# Targeted enrichment outperforms other enrichment techniques and enables more multi-species RNA-Seq analyses

**DOI:** 10.1101/258640

**Authors:** Matthew Chung, Laura Teigen, Hong Liu, Silvia Libro, Amol Shetty, Nikhil Kumar, Xuechu Zhao, Robin E. Bromley, Luke J. Tallon, Lisa Sadzewicz, Claire M. Fraser, David A. Rasko, Scott G. Filler, Jeremy M. Foster, Michelle L. Michalski, Vincent M. Bruno, Julie C. Dunning Hotopp

## Abstract

Enrichment methodologies enable analysis of minor members in multi-species transcriptomic analyses. We compared standard enrichment of bacterial and eukaryotic mRNA to targeted enrichment with Agilent SureSelect (AgSS) capture for *Brugia malayi*, *Aspergillus fumigatus*, and the **Wolbachia** endosymbiont of *B. malayi* (wBm). Without introducing significant systematic bias, the AgSS quantitatively enriched samples, resulting in more reads mapping to the target organism. The AgSS-enriched libraries consistently had a positive linear correlation with its unenriched counterpart (r^2^=0.559-0.867). Up to a 2,242-fold enrichment of RNA from the target organism was obtained following a power law (r^2^=0.90), with the greatest fold enrichment achieved in samples with the largest ratio difference between the major and minor members. While using a single total library for prokaryote and eukaryote in a single sample could be beneficial for samples where RNA is limiting, we observed a decrease in reads mapping to protein coding genes and an increase of multi-mapping reads to rRNAs in AgSS enrichments from eukaryotic total RNA libraries as opposed to eukaryotic poly(A)-enriched libraries. Our results support a recommendation of using Agilent SureSelect targeted enrichment on poly(A)-enriched libraries for eukaryotic captures and total RNA libraries for prokaryotic captures to increase the robustness of multi-species transcriptomic studies.

## Introduction

Dual species transcriptomic experiments have been increasingly employed as a method to analyze the transcriptomes of multiple species within a system[1–9]. However, obtaining a sufficient quantity of reads from each organism is a major issue when simultaneously analyzing the transcriptomes of multiple organisms within a single sample. When extracting total RNA from a host sample, the signal from host transcripts typically overwhelms the signal from secondary organism transcripts under most biologically meaningful conditions. Differential enrichments have been designed to physically extract RNA from secondary organisms in samples by depleting the highly abundant rRNAs and selecting transcripts based on differing properties between the two organisms, such as the differential poly-adenylation status of transcripts [10]. However, in eukaryote-eukaryote dual species transcriptomics experiments, these differences are often not present. Additionally, methods used to enrich for prokaryotic RNA from samples dominated by eukaryotic RNA can fail when the poly(A)-depletion method is unable to remove eukaryotic long non-coding RNAs that lack 3‘-polyadenylatation, and when the efficacy of rRNA depletion is low, as observed with some organisms.

The Agilent SureSelect platform serves as a hybridization-based enrichment method that uses specific baits to target transcripts of interest (www.genomics.agilent.com). By designing sequence-specific baits for an organism, it becomes possible to extract transcripts of interest, such as mRNAs, while avoiding unwanted RNA types, such as rRNAs and tRNAs. In dual species transcriptomics, one of the main advantages of the Agilent SureSelect system is the ability to extract reads in systems where an organism has low relative abundance [11]. To test the efficacy of the Agilent SureSelect platform on multiple dual-and tripartite-species systems where insufficient numbers of reads are obtained from the minor member(s), we designed Agilent SureSelect baits for: (1) *Brugia malayi*, a filarial nematode and the causative agent of lymphatic filariasis, (2) wBm, the obligate mutualistic Wolbachia endosymbiont of B.malayi and (3) *Aspergillus fumigatus* AF293, a fungal pathogen known to cause aspergillosis in immunocompromised individuals.

In filarial nematode transcriptomics, it is relatively straightforward to isolate nematode samples from the mammalian/definitive host while minimizing contaminating host material. Therefore, the Brugia transcriptome can be easily obtained from poly(A)-enriched RNA from Brugia samples originating from the mammalian definitive host, which is often an experimentally infected gerbil (Meriones unguiculatus). However, sampling the Wolbachia endosymbiont transcriptome in these worms is problematic because the endosymbionts contribute less RNA to the total RNA pool. Therefore, wBm reads are overwhelmed by *B. malayi* reads and must be enriched using a rRNA- and poly(A)-depletion. In the mosquito vector host, the parasitic worms develop in the thoracic muscles where they are not easily isolated. Therefore, whole infected mosquito thoraces containing the larval *B. malayi* are used for RNA isolation. As such the *B. malayi* reads are overwhelmed by reads from those of the mosquito vector, typically experimentally infected Aedes aegypti, and the reads of endogenous wBm are even further dwarfed compared to samples obtained from the definitive host. In a separate dual-species transcriptomics experiment examining A fumigatus infections, *A. fumigatus* reads are overwhelmed by reads from the human or mouse host making it difficult to analyze the fungal transcriptome in this host-pathogen interaction. For each eukaryotic system, the performance of the poly(A)-enrichment was compared to that of poly(A)-enrichment supplemented with the Agilent SureSelect platform in extracting *B. malayi* and *A. fumigatus* reads. Additionally, the efficacy of the Agilent SureSelect platform in extracting prokaryotic reads was determined by comparing the total RNA Agilent SureSelect capture to the RiboZero-treated, poly(A)-depletion method in extracting wBm reads.

## Methods

### Mosquito preparation

Aedes aegypti black-eyed Liverpool strain mosquitoes were obtained from the NIH/NIAID Filariasis Research Reagent Resource Center (FR3) and maintained in the biosafety level 2 insectary at the University of Wisconsin Oshkosh (UWO). Desiccated mosquito eggs were hatched in deoxygenated water and the larvae maintained on a slurry of ground TetraMin fish food (Blacksburg, VA, USA) at 27 °C and 80% relative humidity. Female pupae were separated from males using a commercial larval pupal separator (The John Hock Company, Gainesville, FL, USA) and maintained on cotton pads soaked in sucrose solution. Adult female mosquitos were deprived of sucrose ˜8 h prior to blood feeding. Mosquitoes were infected with the *B. malayi* FR3 strain by feeding on microfilaremic cat blood (FR3) through parafilm via a glass jacketed artificial membrane feeder. Microfilaremic cat blood was diluted with uninfected rabbit blood to achieve a suitable parasite density for infection (100-250 mf (microfilariae)/20 µL) and mosquitoes were allowed to feed to repletion. Mosquitoes were maintained in insect incubators until time of worm harvest.

### Nematode preparation

To examine larval development in the vector, groups of mosquitoes were sampled at 18 h post infection (hpi), 4 days post infection (dpi), and 8 dpi. Because larval development occurs in the thoracic muscle of the mosquito, thoraces of infected mosquitoes containing larval *B. malayi* were separated from the head, abdomen, legs, and wings; flash frozen in liquid nitrogen; and stored at -80 °C prior to RNA isolation. To generate third stage larvae (L3) of *Brugia malayi*, infected mosquitoes at 9-16 dpi were processed in bulk using the NIAID/NIH Filariasis Research Reagent Resource Center (FR3) Research Protocol 8.4 (www.filariasiscenter.org). Larvae were isolated in RPMI media containing 0.4 U penicillin and 4 µg streptomycin per mL (RPMI + P/S), flash frozen in liquid nitrogen, and stored at -80 °C. To generate fourth larval stages (L4) and adult worms, freshly isolated L3s were injected into the peritoneal cavities of Mongolian gerbils (Meriones unguiculatus). Briefly, male Mongolian gerbils three months of age or older (Charles River, Wilmington, MA, USA) were anesthetized with 5% isoflurane, immobilized on a thermal support, and administered ocular Paralube. The inguinal region for each gerbil was shaved and disinfected with iodine. Infections were performed by delivering L3 larvae into the peritoneal cavity using a butterfly catheter, which was left in place and flushed afterwards with 1 mL warmed RPMI + P/S to ensure delivery of all larvae. Afterwards, gerbils were removed from the plane of anesthesia and allowed to fully recover in a hospital cage prior to returning to group housing. Worms were later harvested by euthanizing the gerbils and soaking the peritoneal cavities in warm RPMI + P/S. Worms were washed in warm RPMI +P/S to remove traces of gerbil tissue and flash frozen with liquid nitrogen and stored at -80 °C. All animal care and use protocols were carried out in accordance with the relevant guidelines and regulations and were approved by the UWO IACUC.

### Mosquito/Nematode/*Wolbachia* RNA isolation

Mosquito thoraces were combined with TRIzol (Zymo Research, Irvine, CA, USA) at a ratio of 1 mL TRIzol per 50-100 mg mosquito tissue while nematode samples were processed using a 3:1 volume ratio of TRIzol to sample. β-mercaptoethanol was added to a final concentration of 0.1%. The tissues were homogenized in a TissueLyser (Qiagen, Germantown, MD) at 50 Hz for 5 min. The homogenate was transferred to a new tube and centrifuged at 12,000 × g for 10 min at 4 °C. After incubating at room temperature for 5 min, 0.2 volumes of chloroform were added. The samples were shaken by hand for 15 s, incubated at room temperature for 3 min, then loaded into a pre-spun, phase lock gel heavy tube (5Prime, Gaithersburg, MD, USA) and centrifuged for 5 min at 12,000 × g at 4 °C. The upper phase was removed to a new tube and one volume of 100% ethanol was added prior to loading onto a PureLink RNA Mini column (Ambion, Austin, TX). The samples were then processed following manufacturer instructions, quantified using a Qubit fluorometer (Qiagen, Germantown, MD, USA) and/or a NanoDrop spectrometer (NanoDrop, Wilmington, DE, USA). The RNA was then shipped on dry ice to the University of Maryland where it was treated with the TURBO DNA-free kit (Ambion, ThermoFisher Scientific, Waltham, MA, USA) according to the manufacturer‘s protocol.

### **A. fumigatus** infection

Two different models of invasive aspergillosis, both using male BALB/c mice, were employed [12, 13]. In the non-neutropenic model, the mice were immunosuppressed with 5 doses of cortisone acetate, with 500 mg/kg administered subcutaneously every other day four days pre-infection. In the leukopenic model, the mice were administered cyclophosphamide, 250 mg/kg intraperitoneally, and cortisone acetate, 250 mg/kg subcutaneously, on two days pre-infection. On day 4 after infection, the mice were given a second dose of cyclophosphamide, 200 mg/kg intraperitoneally, and cortisone acetate, 250 mg/kg subcutaneously. All mice were infected by placing them in a chamber containing an aerosol of *A. fumigatus* conidia for 1 h. On days 2 and 4 for the non-neutropenic model and on days 4 and 7 for the leukopenic model, 3 mice per time point were sacrificed, after which their lungs were harvested and stored in RNAlater (Ambion, ThermoFisher Scientific, Waltham, MA, USA) for subsequent RNA isolation.

### **A. fumigatus** RNA isolation

Each lung sample was placed in a lysing matrix C tube (MP Biomedicals, Santa Ana, CA, USA) with a single 0.25 inch diameter ceramic sphere (MP Biomedicals, Santa Ana, CA, USA) and homogenized with a bead beater (FastPrep FP120, Qbiogene, Montreal, Quebec, Canada).The RNA was isolated using the RiboPure kit (Ambion, ThermoFisher Scientific, Waltham, MA, USA) following the manufacturer‘s instructions, treated with DNase I (ThermoFisher Scientific, Waltham, MA, USA), and purified using the RNA clean & concentrator kit (Zymo Research, Irvine, CA, USA).

### Illumina NEBNext Ultra Directional RNA libraries without capture

Whole transcriptome libraries were constructed for sequencing on the Illumina platform using the NEBNext Ultra Directional RNA Library Prep Kit (New England Biolabs, Ipswich, MA, USA). When targeting eukaryotic mRNA, polyadenylated RNA was isolated using the NEBNext Poly(A) mRNA magnetic isolation module. When targeting bacterial mRNA, samples underwent rRNA- and poly(A)-reductions, as previously described [10, 14]. SPRIselect reagent (Beckman Coulter Genomics, Danvers, MA, USA) was used to purify cDNA between enzymatic reactions and perform size selection. For indexing, the PCR amplification step was performed with primers containing a 7-nt index sequence. Libraries were evaluated using the GX touch capillary electrophoresis system (Perkin Elmer, Waltham, MA) and sequenced on a HiSeq2500 generating 100-bp paired end reads.

### *Wolbachia* custom RNA capture

Pre-capture libraries were constructed from 500-1000 ng of total RNA samples using NEBNext Ultra Directional RNA Library Prep kit (NEB, Ipswich, MA, USA). First strand cDNA was synthesized without mRNA extraction to retain non-polyadenylated transcripts and was fragmented at 94°C for 8 min. After adaptor ligation, cDNA fragments were amplified with 10 cycles of PCR before capture. Wolbachia transcripts were captured from 200 ng of the amplified libraries using an Agilent SureSelectXT RNA (0.5-2 Mbp) bait library designed specifically for wBm. Library-bait hybridization reactions were incubated at 65 C for 24 h then bound to MyOne Streptavidin T1 dynabeads (Invitrogen, Carlsbad, CA, USA). After multiple washes, bead-bound captured library fragments were amplified with 18 cycles of PCR. The libraries were loaded on a HiSeq4000 generating 151-bp paired end reads.

### **B. malayi** custom RNA capture

Pre-capture libraries were constructed from 1000 ng of total RNA samples using NEBNext Ultra Directional RNA Library Prep kit (NEB #E7420, Ipswich, MA, USA). Except where noted, poly(A)-enrichment according to the manufacturer‘s protocol was used. In comparisons of capture on poly(A)-enriched libraries with total RNA libraries, total RNA libraries were constructed after first strand cDNA was synthesized without mRNA extraction to retain non-polyadenylated transcripts and fragmented at 94°C for 8 min. After adaptor ligation, cDNA fragments were amplified for 10 cycles of PCR before capture. *B. malayi* transcripts were captured from 200 ng of the amplified libraries using an Agilent SureSelectXT Custom (12-24 Mbp) bait library designed specifically for *B. malayi*. Library-bait hybridization reactions were incubated at 65°C for 24 h then bound to MyOne Streptavidin T1 Dynabeads (Invitrogen, Carlsbad, CA). After multiple rounds of washes, bead-bound captured library fragments were amplified with 16 cycles of PCR. The libraries were loaded on a HiSeq4000 generating 151-bp paired end reads.

### *Aspergillus* custom RNA capture

Pre-capture libraries were constructed from 700 ng of total RNA samples using NEBNext Ultra Directional RNA Library Prep kit (NEB #E7420, Ipswich, MA, USA) according to the manufacturer‘s protocol. Briefly, mRNA was extracted with oligo-d(T) beads and reverse-transcribed into first strand cDNA with random primers. First strand cDNA was fragmented at 94°C for 8 min. After adaptor ligation, cDNA fragments were amplified in thermocycler for 10 cycles before capture. Aspergillus transcripts were captured from 100 ng of amplified libraries Agilent SureSelectXT Custom (12-24 Mbp) bait library for *A. fumigatus*. Libraries were incubated at 65°C for 24 h then bound to MyOne Streptavidin T1 Dynabeads (Invitrogen, Carlsbad, CA, USA). After multiple rounds of washes, bead-bound captured library fragments were amplified with 16 cycles of PCR. The libraries were loaded on a HiSeq4000 generating 151-bp paired end reads.

### Sequence alignment, feature counts, and fold enrichment calculations

The sequence reads originating from all samples used in all *B. malayi* and *A. fumigatus* AF293 comparisons were mapped to the *B. malayi* genome WS259 (www.wormbase.org) or *A. fumigatus* AF293 genome CADRE34 (www.aspergillusgenome.org), respectively, using the TopHat v1.4 [15] aligner. Read counts for all genes were obtained using HTSeq (v0.5.3p9) [16], with the mode set to “union.“ For the **B. malayi** genome, RNAmmer v1.2 [17] identified 5 protein-coding genes (WBGene00228061, WBGene00268654, WBGene00268655, WBGene00268656, WBGene00268657) overlapping with predicted rRNAs. Reads overlapping with these genes were excluded from all gene-based analyses. The sequencing reads for the wBm comparisons were mapped to the wBm assembly [18] using Bowtie v0.12.9 [19]. Counts for each wBm gene were calculated by summing the sequencing depth per base pair, obtained using the DEPTH function of SAMtools v1.1 [20], for each of the unique genomic positions in each gene. This value was then divided by the average read length for the sample, yielding a read count value for each gene. Across all comparisons, read counts were normalized between each of the different samples through a conversion to transcripts per million (TPM).

The fold enrichment, *F*_*e*_, of reads belonging to an organism conferred by the Agilent SureSelect compared to the other enrichment techniques was calculated as

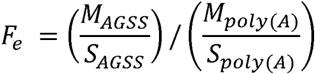

the ratio of the number of reads mapped with Agilent SureSelect capture (*M_AGSS_*) to the number of reads sequenced with Agilent SureSelect capture (*M_AGSS_*) divided by the ratio of number of reads mapped with poly(A)-enrichment or the rRNA-, poly(A)-depletion (*M_poly(A)_*) to the number of reads sequenced with poly(A)-enrichemnt or the rRNA-, poly(A)-depletion (*S_poly(A)_*. By taking a ratio of the percentage of mapped reads for each enrichment technique, the fold enrichment value represents an amount of enrichment conferred by the Agilent SureSelect when compared to the other enrichment technique for a given sample. The fold enrichment values when comparing the efficacy of Agilent SureSelect capture on poly(A)-enriched libraries to total RNA libraries were calculated similarly as

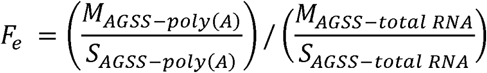

where *M_AGSS-poly(A)_* and *S_AGSS-poly(A)_* represent the mapped and sequenced reads from poly(A)-enriched Agilent SureSelect capture libraries, while *M_AGSS-total RNA_* and *S_AGSS-total RNA_* represent the mapped and sequencing reads from Agilent SureSelect capture from the total RNA libraries.

## Results

### Comparison of poly(A)-enriched Agilent SureSelect to only poly(A)-enrichment for *B. malayi*

Poly(A)-enriched, Agilent SureSelect libraries were compared to only poly(A)-enriched libraries from six RNA samples originating from three different time points of the *B. malayi* vector life stages (18 hpi, 4 dpi, and 8 dpi) (**Figure 1a**). Across these six samples, the poly(A)-enrichment alone yielded 0.38-23.35% of reads mapping to the *B. malayi* genome, with 61.25-82.51% of reads mapping to the A. aegypti genome. Of the poly(A)-enriched libraries capture using the Agilent SureSelect baits, 56.14%-81.52% of reads mapped to the *B. malayi* genome, with 2.28-24.02% of reads mapping to the A. aegypti genome (**Supplementary Table 1**). The fold enrichment conferred to libraries captured with Agilent SureSelect was found to be inversely proportional to the number of reads mapped to its poly(A)-enrichment only counterpart, such that experiments with the fewest reads mapped in the poly(A)-enrichment had the highest fold enrichment with the Agilent SureSelect capture. The 18 hpi samples, which have the lowest percentage of mapped reads to the *B. malayi* genome has the greatest fold enrichment (110-146x) while the 8 dpi samples, with the highest percentage of *B. malayi* mapped reads, has the lowest fold enrichment (3-4x).

**Figure 1.**
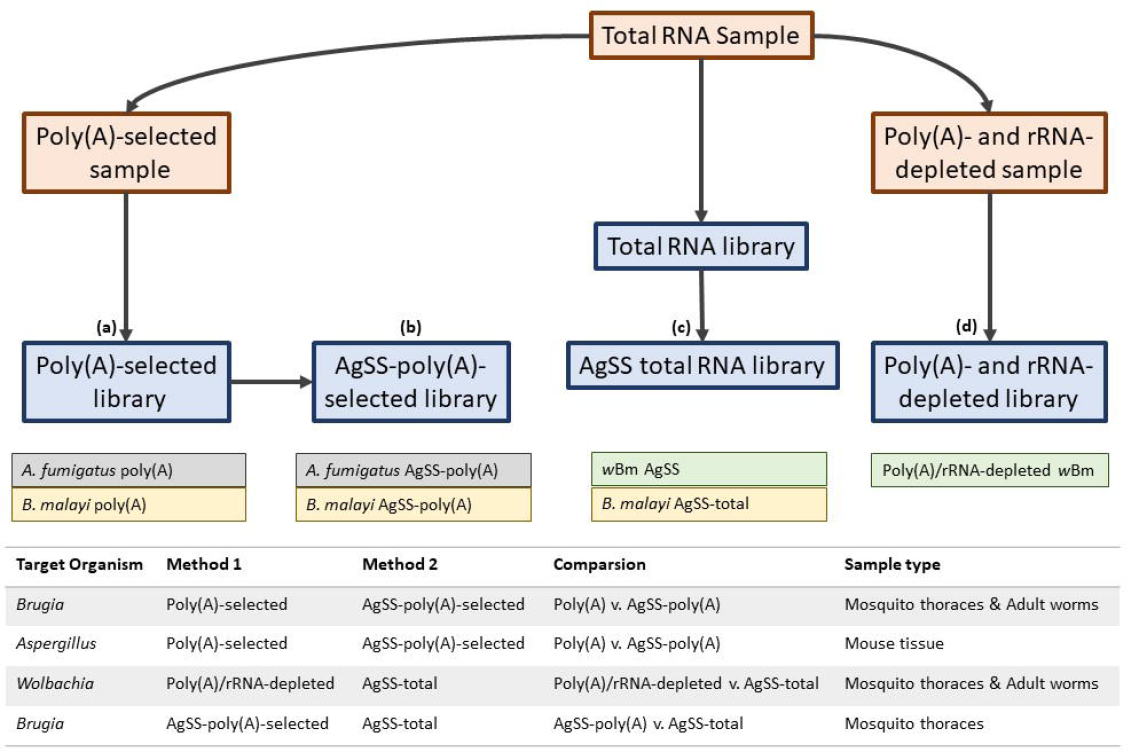
Sample and library preparation. This schematic illustrates the sample (auburn rectangle) and library (blue rectangle) preparation workflow terminating in the libraries that were loaded on the Illumina sequencer. (a) For *B. malayi* and *A. fumigatus*, a poly(A)-selected sample was created from an aliquot of total RNA that was used to create a poly(A)-selected library. (b) The *B. malayi* or *A. fumigatus* Agilent SureSelect (AgSS) baits were subsequently used to capture the targeted RNA from poly(A)-selected libraries. (c) For AgSS-enriched wBm libraries, an RNA library was constructed from an aliquot of total RNA that underwent targeted enrichment with the Wolbachia AgSS baits. Unlike the eukaryotic enrichments, the bacterial AgSS capture is performed on total RNA. For a limited number of libraries described in the text, an RNA library was constructed from an aliquot of total RNA (i.e. without poly(A)-enrichment) that underwent targeted enrichment with the Brugia AgSS baits. (d) For poly(A)/rRNA-depleted libraries enriched for wBm, an aliquot of total RNA from either mosquito thoraces or adult nematodes was enriched for bacterial mRNA by removing Gram-negative and human rRNAs with two RiboZero removal kits and polyadenylated RNAs with DynaBeads.

A principal component analysis (PCA) using the log_2_-TPM values for the 11,085 predicted *B. malayi* protein-coding genes reveals that the *B. malayi* samples cluster based on life stage rather than library preparation (**Figure 2a**), suggesting that the capture does not introduce a systematic bias to the transcriptome of a sample. The exceptions are the 18 hpi vector poly(A)-enrichment-only samples, which we attribute to these samples having an insufficient number of reads mapped to features (143,078 and 212,342 reads, accounting for <20 reads per gene assuming equal distribution) to accurately represent the transcriptome. Supporting this, in the 18 hpi samples, an average of 2,460 protein coding genes (22.18% of the *B. malayi* protein coding genes) had reads mapping to them in the poly(A)-enriched Agilent SureSelect samples but not its poly(A)-enriched counterpart, while in the 4 dpi and 8 dpi samples, an average of 952 (8.58%) and 566 (5.09%) genes, respectively, had reads mapping to them from the poly(A)-enriched Agilent SureSelect libraries but not the poly(A)-enriched libraries. Collectively, across all 6 samples, 392 unique genes were detected in only the poly(A)-enrichment, but not the poly(A)-enriched Agilent SureSelect libraries in at least one comparison. The Agilent SureSelect probe design was based on an older version of the annotation compared to the version used for feature calling, and as such, 91 of these 392 genes were not covered by a probe.

**Figure 2.**
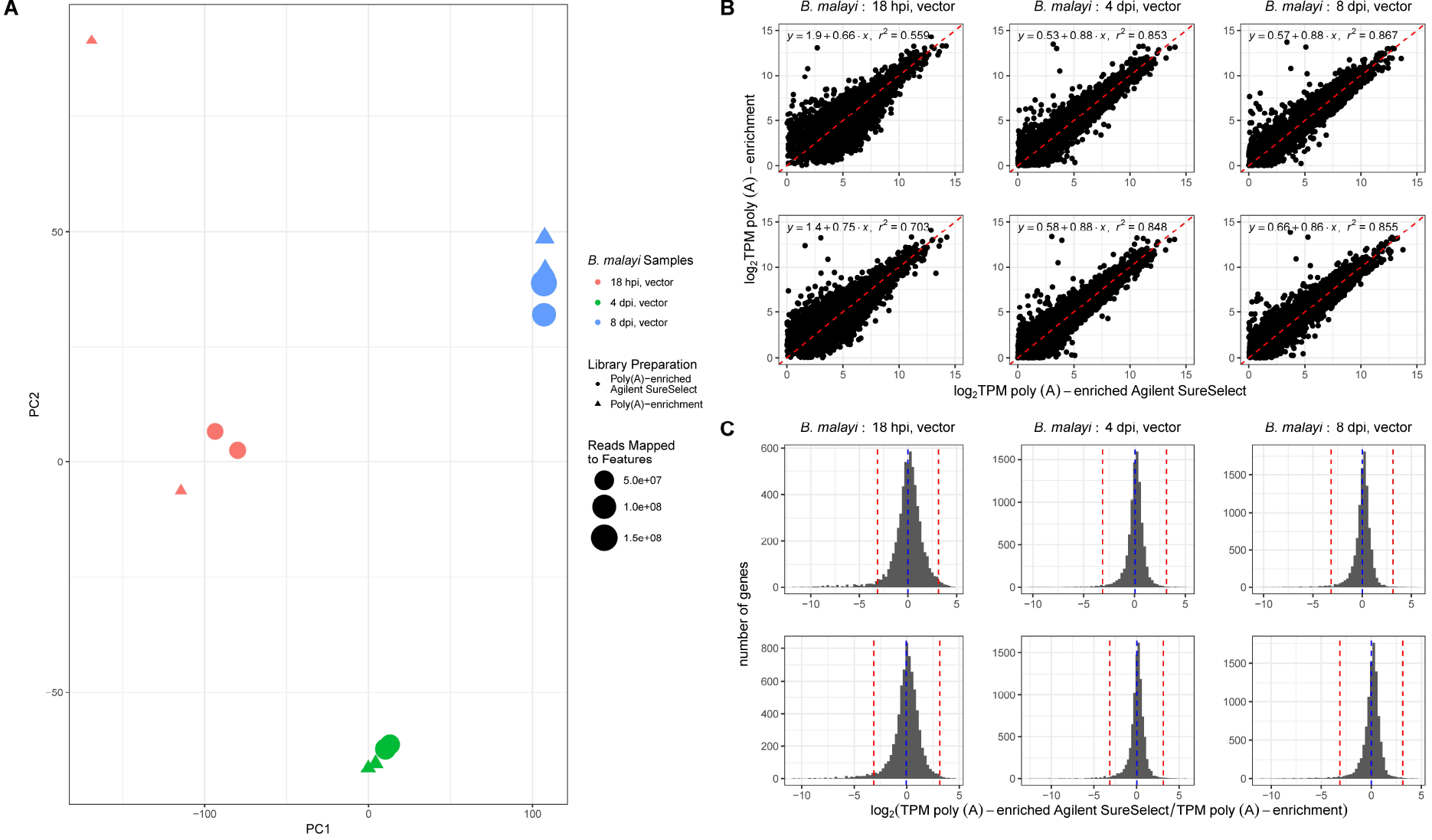
Comparison of the poly(A)-enriched to the poly(A)-enriched Agilent SureSelect **B. malayi** transcriptomes from 18 hpi, 4 dpi, and 8 dpi vector life stages. (a) A principal component analysis (PCA) plot was generated using the log_2_ TPM values for the poly(A)-enriched only (circles) and poly(A)-enriched Agilent SureSelect (triangles) *B. malayi* from 18 hpi (red), 4 dpi (green), and 8 dpi (blue) vector samples. The size of each symbol denotes the number of reads mapped to protein-coding genes for each sample. The samples cluster based on *B. malayi* sample rather than by the enrichment method, indicating the poly(A)-enriched Agilent SureSelect does not substantially differ from its poly(A)-enrichment only counterpart in representing the *B. malayi* transcriptome for the shown samples. Samples with the lowest number of reads (e.g. the 18 hpi samples) cluster further from their replicates than expected, likely attributed to an insufficient number of reads and indicating that enrichment is most desirable in these situations. (b) For each sample, the log_2_ TPM values for genes in the poly(A)-enriched Agilent SureSelect samples were plotted against the log_2_ TPM values for genes in the poly(A)-enrichment only samples when expression was observed with both enrichment methods for the sample. Genes with similar expression in the Agilent SureSelect and the poly(A)-enrichment samples are expected to lie close to the identity line (x=y; red). Genes whose expression values are more elevated in the poly(A)-enriched Agilent SureSelect sample compared to the poly(A)-enrichment sample lie below the identity line while genes more elevated in the poly(A)-enrichment compared to the poly(A)-enriched Agilent SureSelect lie above the identity line. (c) For all genes expressed using both enrichment methods for each sample, the frequency distribution of the log_2-_transformed ratio of the poly(A)-enriched Agilent SureSelect TPM to the poly(A) enrichment TPM was plotted. Genes with log_2_ ratio values >0 have higher expression values in the poly(A)-enriched Agilent SureSelect sample while log_2_ ratio values <0 are representative of genes with higher expression values with the poly(A) enrichment. Significantly biased genes were defined as those differentiating by >3 standard deviations (red) from the mean of the log_2_-transformed ratio values (red). Across each comparison, the number of genes with significantly elevated expression in the Agilent SureSelect comprised at most 0.31% of the total number of *B. malayi* protein-coding genes while the number of genes with significantly elevated expression in the poly(A)-enrichment samples comprised at most 1.52% of *B. malayi* protein-coding genes.

The log_2_ TPM values from the poly(A)-enriched Agilent SureSelect and the poly(A)-enriched libraries had a positive linear correlation with r^2^ values ranging from 0.559-0.867 (**Figure 2b**). The 4 dpi and 8 dpi correlations were the strongest (**Figure 2b**), for the reasons discussed above, namely that fewer reads were recovered from the 18 hpi vector samples without Agilent SureSelect enrichment. To identify individual genes in which the library preparations affected the calculated TPM, the log_2_ ratio of the poly(A)-enriched Agilent SureSelect TPM relative to the poly(A) enrichment TPM for each gene was calculated (**Figure 2c**). Genes with relatively similar expression levels in the two library preparations would have a log_2_ ratio of 0 while genes more highly expressed in the poly(A)-enriched Agilent SureSelect preparation would be >0 and genes more highly expressed in the poly(A) enrichment would be <0. Genes with a log_2_-ratio value of >3 standard deviations from the mean were defined as having significantly altered levels of expression between the two libraries. Across all six comparisons, the number of genes with significantly higher TPM values in the Agilent SureSelect libraries never accounted for >0.31% (<34 genes) of all protein-coding genes in the *B. malayi* genome. The number of genes with significantly higher TPM values in the poly(A)-enrichment ranged from 0.93%-1.52% (103-168 genes) of all protein-coding genes in *B. malayi* across the six comparisons (Supplementary Table 1).

### Comparison of poly(A)-enriched Agilent SureSelect and only poly(A)-enrichment for *A. fumigatus*

The *A. fumigatus* AF293 transcriptome from poly(A)-enriched Agilent SureSelect and poly(A)-enriched only libraries were compared for 11 RNA samples from immunocompromised mice or neutropenic mice infected with *A. fumigatus* AF293 (**Figure 1a**). Of the 11 samples, 6 samples originated from three replicates of immunocompromised mice taken 2 dpi and 4 dpi with *A. fumigatus*. The remaining 5 samples consist of two replicates of neutropenic mice taken 4 dpi with *A. fumigatus* and three replicates of neutropenic mice taken 7 dpi with *A. fumigatus*.

Across all 11 samples, the total percentage of reads mapping to *A. fumigatus* from the poly(A)-enriched samples was consistently ≤0.02%, while 0.30%-11.74% of reads mapped in the poly(A)-enriched Agilent SureSelect samples. In the poly(A)-enrichment only samples, a maximum of 27,531 reads mapped to *A. fumigatus* AF293, accounting for <3 reads/gene if they were evenly distributed. As such, comparisons between Agilent SureSelect and poly(A)-enriched libraries are not possible.

There are 4,168-7,387 genes (41.68%-76.70%) that are identified only in the poly(A)-enriched Agilent SureSelect libraries but not the poly(A)-enriched libraries, compared to the maximum of 52 genes identified only in the poly(A)-enriched libraries across all *A. fumigatus* comparisons (**Supplementary Table 2**). For all 11 comparisons, using the Agilent SureSelect methodology conferred a 614x to 1,212x fold enrichment of *A. fumigatus* AF293 reads relative to the poly(A)-enrichment, demonstrating the power of this technology in dual-species transcriptomics, going from data that was unamenable to a traditional transcriptomics analysis to obtaining data where robust statistical tests can be performed. The proportion of reads mapping to features relative to the total number of reads mapped is consistent between samples of the same library preparation, with *A. fumigatus* poly(A)-enriched Agilent SureSelect samples ranging from 52.23-54.87% and the poly(A)-enrichment samples ranging from 45.89-51.22%.

### Comparison of total RNA Agilent SureSelect to rRNA-, poly(A)-depleted for the bacterial *Wolbachia* endosymbiont *w*Bm

With both of the previously discussed biological systems, we tested the effectiveness of the Agilent SureSelect platform to study eukaryote-eukaryote host-pathogen interactions. By enriching for wBm reads in *B. malayi* samples, we sought to test the effectiveness of Agilent SureSelect libraries in extracting prokaryotic reads. The standard protocol for Agilent SureSelect capture in eukaryotes relies on capturing targets from poly(A)-enriched libraries, but for prokaryotic samples the capture is performed on total RNA libraries. These total RNA Agilent SureSelect libraries were compared to rRNA-, poly(A)-depleted library preparations from nine *B. malayi* samples taken at various stages of the life cycle to analyze the transcriptome of its Wolbachia endosymbiont, wBm (**Figure 1b**). Three RNA samples were obtained from the mammalian portion of the Brugia lifecycle where worms are large enough to be physically separated from the mammalian tissue. These RNA samples generate predominantly Brugia reads with a minority of Wolbachia reads. Of these three samples, one sample was recovered at 24 dpi, representative of female worms just prior to the final L4 to adult molt, while the other two samples are from two reproductively mature adult females recovered 7 months post-infection. The other six samples are from the vector portion of the *B. malayi* lifecycle where tripartite RNA samples are obtained, consisting of predominantly mosquito reads, with a smaller percentage of Brugia reads, and an even smaller percentage of Wolbachia reads. These six samples consisted of two A. aegypti replicates each of the 18 hpi, 4 dpi, and 8 dpi with *B. malayi*, representative of the L1 through early L3 *B. malayi* life stages. Of the six samples taken from the vector life stages, at most 1,552 reads (<0.01% of sequenced reads) mapped to the wBm genome from the rRNA-, poly(A)-depleted samples, of which 729 reads were identified as mapping to protein-coding genes for an average of <1 read mapped/gene (Supplementary Table 3). In contrast, the rRNA-, poly(A) depleted adult female *B. malayi* samples taken from the mammalian host had a minimum of 17,107 reads mapping to wBm protein-coding genes, equating to ~20 reads/gene while the 24 dpi sample had 246,549 reads mapped to protein-coding genes, equating to ~294 reads/gene.

The Agilent SureSelect-enriched samples had an increased number of wBm-mapping reads for all 9 samples, with 20.4-37.4 million reads (18.68%-32.53%) mapping to wBm genes for the three gerbil samples and 0.2-4.0 million reads (0.46-6.43% of sequenced reads) mapping to wBm genes for the mosquito vector samples. The total RNA Agilent SureSelect libraries from gerbil samples were 20-41x fold enriched for wBm reads relative to the rRNA-, poly(A)-depleted libraries while the libraries from vector samples were 353-2,242x fold enriched. The proportion of wBm reads mapped to protein-coding genes were consistently higher in the Agilent SureSelect in both the gerbil and vector samples, ranging from 88.31-92.44% and 44.34-56.14%, respectively, compared to the 10.62-20.11% and 45.21-58.73% observed in the rRNA-, poly(A)-depleted samples. This difference in the number of wBm reads mapping to protein-coding genes can be partially attributed to a greater number of reads mapping to rRNAs in the rRNA-, poly(A)-depleted samples, with 18.73-33.82% of wBm mapped reads mapping to rRNAs compared to 0.1-2.1% observed in the total RNA, Agilent SureSelect samples (Supplementary Table 3). A principal component analysis using the TPM values of the three gerbil life stage comparisons demonstrates that individual samples segregate based on library preparation (Figure 3a). All three of the total RNA Agilent SureSelect samples are clustered close together, while the rRNA-, poly(A)-depleted samples are further distributed, likely due to the insufficient coverage of the wBm transcriptome in the absence of Agilent SureSelect enrichment. The mosquito vector life stage samples were not used for the principal component analysis due to an insufficient number of reads mapping in the poly(A)-depletion portion of the comparison to compare the two enrichment methods. Like the *B. malayi* comparisons, the log_2_ TPM values from the wBm Agilent SureSelect libraries and the rRNA- and poly(A)-depletion libraries made from mammalian life stages had a positive linear correlation with r^2^ values ranging from 0.575-0.744 (Figure 3b).

**Figure 3.**
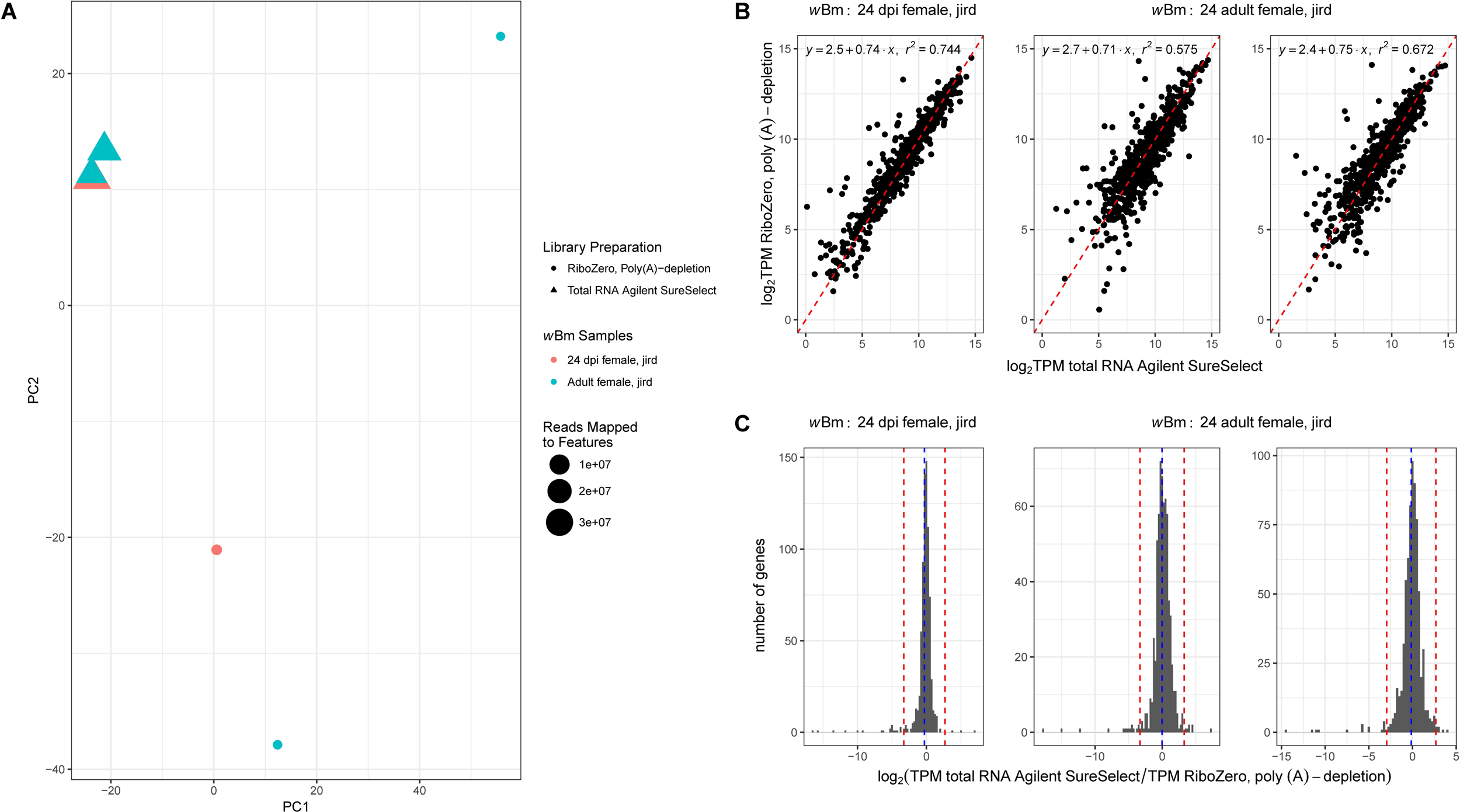
Comparison of the rRNA-, poly(A)-depleted to the total RNA Agilent Sure Selectw Bm transcriptomes from the female 24 dpi and adult *B. malayi*life stages. (a) A principal component analysis of the rRNA-, poly(A)-depleted (triangle) and total RNA Agilent SureSelect (circle) wBm transcriptome from three samples of the *B. malayi* gerbil life stage was conducted, with one sample from female 24 dpi *B. malayi* (red) and two samples from adult female *B. malayi* (blue). The size of each point is relative to the number of reads mapped to protein-coding genes for each sample. In this instance, the samples no longer cluster based on the sample type. Instead, the samples distinctly cluster apart based on enrichment method, most likely due to the smaller number of reads present in samples lacking the Agilent SureSelect capture. Consistent with this, the sample with the fewest reads, one of the rRNA-, poly(A)-depleted adult females, is most distant. The three samples enriched using the Agilent SureSelect cluster together, possibly indicating the similarity of wBm transcriptomes originating from female 24 dpi and adult *B. malayi*. (b) For each sample, the log_2_ TPM values for genes detected with the Agilent SureSelect capture on total RNA were plotted against the log_2_ TPM values for genes detected with the rRNA-, poly(A)-depletion when a gene was detected in both enrichments. Genes with similar expression in both the total RNA Agilent SureSelect and the rRNA-, poly(A)-depleted samples are expected to lie close to the identity line (x=y; red). Genes whose expression values are more elevated in the total RNA Agilent SureSelect compared to the rRNA-, poly(A)-depletion lie below the identity line while genes more elevated in the rRNA-, poly(A)-depletion compared to the total RNA Agilent SureSelect lie above the identity line. (c) The frequency distribution log_2_-transformed ratio of the total RNA Agilent SureSelect TPM to the rRNA- and poly(A)-depleted TPM for each gene detected in both enrichment methods was plotted. The blue, dashed line indicates the mean log_2_-transformed ratio while the red dashed line marks two standard deviations from the mean log_2_-transformed ratio. Genes with log_2_ ratio values >2 standard deviations above or below the mean were marked as biased towards the rRNA-, poly(A)-depletion, total RNA Agilent SureSelect or the rRNA-, poly(A)-depletion, respectively. Across all 9 comparisons, an average of ~2.9 genes were detected with significantly higher expression with the total RNA Agilent SureSelect while an average of ~7.1 genes had significantly higher expression in the rRNA-, poly(A)-depleted samples.

A total of 45 unique genes (~5.4% of protein-coding genes in wBm) were detected in at least one of the three gerbil comparisons in the sample treated using the rRNA-, poly(A)-depletion libraries but not its total RNA Agilent SureSelect counterpart. Of these 45 genes, 41 were not covered by any of the Agilent SureSelect probes while the remaining four genes had <11% of their length covered by a probe. To identify genes with significantly greater expression in one enrichment method over the other, a cutoff of two standard deviations from the log_2_ ratio of the Agilent SureSelect TPM to the rRNA-, poly(A)-depleted TPM was used (Figure 3c). Genes with a log_2_ ratio of >2 standard deviations below the average log_2_ ratio were determined to have significantly higher expression in the rRNA-, poly(A)-depletion sample while genes with a log_2_ ratio >2 standard deviations above the average, were identified as having significantly higher expression in the total RNA Agilent SureSelect sample. Across the three comparisons, a total of 30 unique genes had TPM values significantly higher in the rRNA-, poly(A)-depleted sample compared to the Agilent SureSelect sample in at least one comparison. Of these genes, 17 genes were not covered by any probe, six genes had <17% of their length covered by a probe, and the remaining seven genes are covered in the entirety of their length by the probe design. Additionally, there were 152 unique genes (~18.1% of protein-coding genes in wBm) detected in the total RNA, Agilent SureSelect but not the poly(A)-depletion in at least one of the comparisons. Of the 152 genes, 114 had average TPM values in the bottom quartile of the expressed genes across the three Agilent SureSelect samples, highlighting the ability of the Agilent SureSelect preparation to detect low abundance transcripts.

### Comparison of Agilent SureSelect*B. malayi* transcriptome data from total RNA and poly(A)-enriched samples

For eukaryote transcriptome experiments, the Agilent SureSelect platform is typically preceded by construction of a poly(A)-selected library that is then hybridized to the bait probes. The poly(A)-enrichment step serves to remove high abundance, non-mRNA transcripts, such as rRNAs, tRNAs, and other ncRNAs. For bacterial samples, libraries must be constructed on total RNA since bacterial mRNAs lack the poly(A) tails present in eukaryotes. For eukaryote-prokaryote transcriptomic experiments, it may be advantageous to use the Agilent SureSelect platform for both bacteria and eukaryotes on the same single library constructed from total RNA, especially for clinical samples where RNA may be limiting. Therefore, we sought to compare the Agilent SureSelect enrichment on a library constructed from total *B. malayi* RNA with one constructed following poly(A)-enrichment using two samples taken from the mosquito vector portion of the *B. malayi* life stages at 18 hpi and 8 dpi (Figure 1c).For both the 18 hpi and 8 dpi timepoints, most genes yield comparable log_2_ TPM values between the two enrichment methods (Figure 4a) with <1.5% of genes having a bias towards a particular library construction protocol (Figure 4b). However, despite the 8 dpi comparison yielding a similar number of genes detected unique to each enrichment method, in the 18 hpi sample comparison, 1,033 genes (9.3% of all protein-coding genes) were detected only in the Agilent SureSelect, poly(A)-enriched library while 323 (2.91%) were only detected with the Agilent SureSelect, total RNA library (Supplementary Table 4). In both cases, a majority of the genes detected in only one library construction method were within the bottom 20% of expressed genes in the sequencing data collected, with 815 genes (78.9%) in the selection conducted with the total RNA library and 233 genes (72.14%) in the selection conducted with the poly(A)-enriched library. Similarly, in the 8 dpi sample comparison, 2.35% (261 genes) of *B. malayi* protein-coding genes were identified only in the target captured poly(A)-enriched library while 1.71% (189 genes) were identified only in the target captured total RNA library. Again, in both cases, a majority of the genes detected in only one library construction method were within the bottom 20% of expressed genes in the sequencing data collected, with 236 genes (90.42%) in the selection conducted with the poly(A)-enriched RNA library and with 179 genes (94.71%) in the selection conducted with the total RNA library. Based off these results, both Agilent SureSelect methods can detect low abundance transcripts, but the poly(A)-enriched Agilent SureSelect was able to select for a greater number of these transcripts in both comparisons.

**Figure 4.**
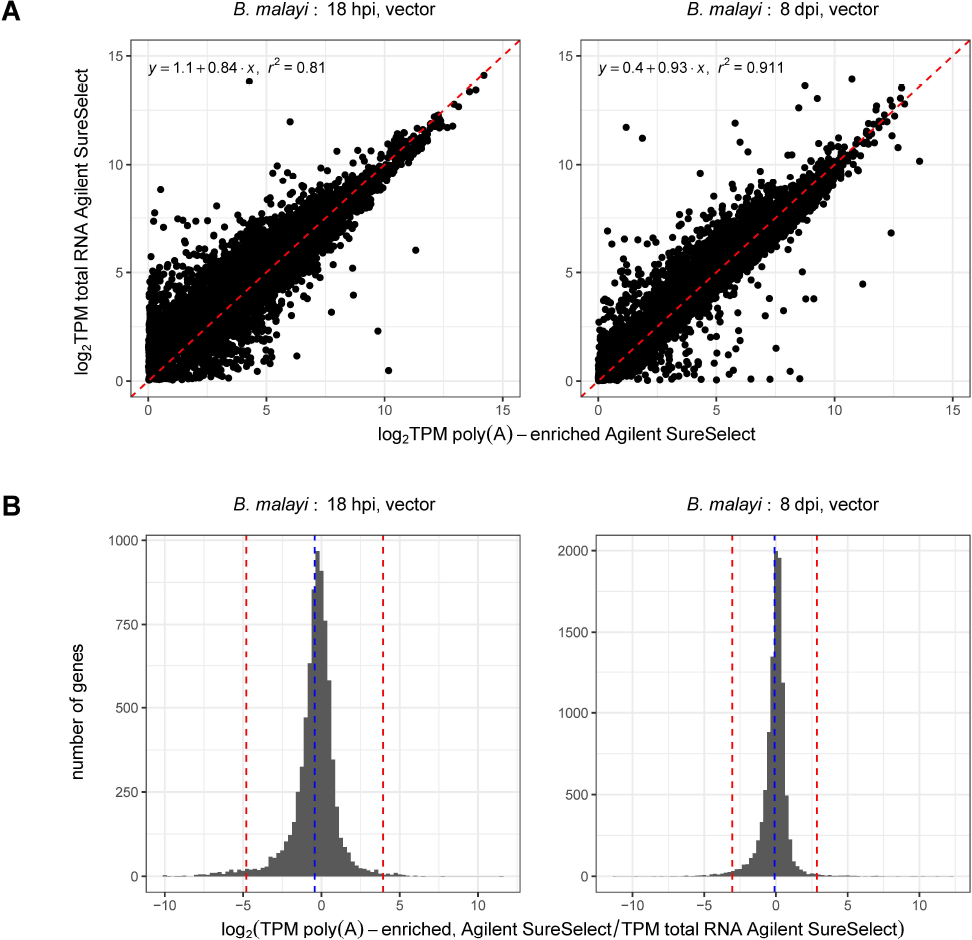
Comparison of the poly(A)-enriched Agilent SureSelect to the total RNA Agilent SureSelect *B. malayi* transcriptomes from 18 hpi and 8 dpi vector life stages. (a) For both the 18 hpi and 8 dpi vector samples, the log_2_ TPM values for each *B. malayi* gene detected in both the poly(A)-enriched Agilent SureSelect and the total RNA Agilent SureSelect were plotted against one another. Most genes in both plots fall along the identity line (y=x; red), indicating their similar expression values in both the poly(A)-enriched Agilent SureSelect and the total RNA Agilent SureSelect libraries. (b) A histogram of the log_2_ ratios of the poly(A)-enriched Agilent SureSelect TPM to the total RNA Agilent SureSelect TPM was generated for each sample using all genes detected with both enrichment methods. The blue dashed line marks the average log_2_ ratio value while the red lines mark three standard deviations from the average log_2_ ratio. Genes with log_2_ ratio values >3 standard deviations above the average are marked as significantly elevated in the poly(A)-enriched Agilent SureSelect while genes with log_2_ ratio values >3 standard deviations below the average are marked as significantly elevated in the total RNA Agilent SureSelect samples. For both the library preparations, ≤1.4% (155 genes) of protein-coding genes are biased towards a single enrichment method, indicating no significant bias towards an enrichment.

The percentage of reads mapping to the *B. malayi* genome were greater in the poly(A)-enriched Agilent SureSelect samples compared to the total RNA Agilent SureSelect samples, with 56.14% and 80.13% of reads mapping to the *B. malayi* genome, compared to 22.96% and 69.84%, of reads mapped for the 18 hpi and 8 dpi samples respectively. The poly(A)-enriched Agilent SureSelect samples were enriched for reads mapping to *B. malayi* by 2.4x for the 18 hpi sample and 1.1x for the 8 dpi sample relative to the total RNA Agilent SureSelect. Additionally, in the 18 hpi and 8 dpi poly(A)-enriched RNA, Agilent SureSelect samples, 46.63% and 46.35% of their mapped reads mapped to protein-coding genes while in the total RNA Agilent SureSelect only 20.22% and 32.21% mapped to protein-coding genes (Supplementary Table 4). This is explained by the increased number of multi-mapping reads observed in the total RNA Agilent SureSelect samples, with 70.29% and 39.29% of the total number of mapped reads in the 18 hpi and 8 dpi vector samples multi-mapping to the *B. malayi* genome. In comparison, only 5.93% and 5.42% of mapped reads were multi-mapping in the 18 hpi and 8 dpi poly(A)-enriched Agilent SureSelect samples. A total of 80.82% and 69.86% of these multi-mapping reads in the total RNA Agilent SureSelect 18 hpi and 8 dpi vector samples were found to be mapping to the *B. malayi* rRNAs compared to only 13.76% and 1.56% in the poly(A)-enriched Agilent SureSelect samples (Supplementary Table 4). The Agilent SureSelect platform on poly(A)-enriched RNA was able to obtain both a higher percentage of *B. malayi* mapped reads and a higher percentage of *B. malayi* reads mapping to protein-coding genes. Therefore, we recommend the use of the Agilent SureSelect enrichment on poly(A)-enriched libraries for eukaryotic systems whenever possible.

## Discussion

With multi-species RNA-Seq experiments emerging at the forefront of experimental designs for transcriptome analyses, novel methods are being developed to more efficiently extract the transcriptomes for multiple organisms from a single sample. By comparing the transcriptomes generated with standard enrichment methods, such as poly(A)-enrichment or rRNA-, poly(A)-depletion, against the Agilent SureSelect platform, we sought to independently validate the use of the Agilent SureSelect in enriching for transcripts originating from a specific organism of interest while minimizing any type of library bias on the transcriptome. When analyzing the *B. malayi* transcriptomes of samples prepared with or without the Agilent SureSelect, most genes could be detected using both library preparations. However, the samples prepared using the Agilent SureSelect enriched the number of reads mapped by 3x-146x compared to samples enriched only with a poly(A)-enrichment and were consistently able to detect a greater number of genes, most of which were low abundance transcripts. Similarly, in the *A. fumigatus* comparisons, the poly(A)-enrichment by itself was unable to extract a sufficient number of *A. fumigatus* reads to accurately represent the transcriptome of the sample. When the poly(A)-enrichment was supplemented with Agilent SureSelect capture, the enrichment of *A. fumigatus* reads was increased 614-1,212x. When using the Agilent SureSelect instead of the rRNA-, poly(A)-depletion preparation for the Wolbachia endosymbiont wBm transcriptome, the Agilent SureSelect samples were enriched for wBm mapped reads by 20-41x in the gerbil life cycle samples and 353-2,242x in the vector life cycle samples.

For each of these samples, if the fold enrichment for the number of mapped reads in the poly(A)-enriched or total RNA Agilent SureSelect is plotted against the percentage of reads mapped in its poly(A)-enrichment or poly(A), rRNA depletion counterpart, the data follow a power law (Figure 5). As expected, the fold enrichment obtained with the Agilent SureSelect capture is inversely proportional to the percentage of mapped reads obtained using the currently used enrichment methods of poly(A)-20 enrichment to capture eukaryotic RNA or rRNA- and poly(A)-depletion to capture bacterial RNA from multi-species samples, indicating samples with a lower relative abundance of RNA from the target organism yield the greatest fold enrichment.

**Figure 5.**
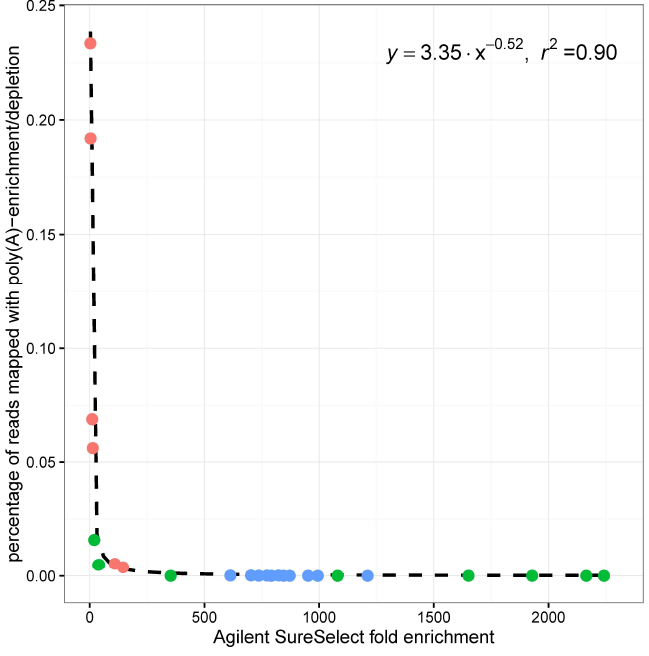
Scatter plot of the percentage of poly(A)-enrichment or depletion reads mapped against the fold enrichment value observed using the Agilent SureSelect follows a power law. The fold enrichment value for each sample used in the poly(A)-enrichment/depletion versus Agilent SureSelect comparisons for *B. malayi* (red), *A. fumigatus*(blue), and wBm (green) was calculated by taking the ratio of the percentage of Agilent SureSelect reads mapped to the percentage of poly(A)-enrichment/depletion reads mapped. The relationship between these fold enrichment values and the percentage of the poly(A)-enrichment/depletion reads mapped to the reference genome can be fitted to a power law. The inversely proportional relationship between the Agilent SureSelect fold enrichment values and the percentage of reads mapped using the percentage of reads mapped using the poly(A)- enrichment/depletion indicates that the fold enrichment conferred using the Agilent SureSelect platform exponentially increases with the exponential decrease of the percentage of reads mapped using the standard enrichment/depletion methods until an ~1300-fold enrichment.

While the Agilent SureSelect can enrich a sample for transcripts of interest, the platform requires probes designed for each gene to avoid any enrichment bias. Most instances of the Agilent SureSelect failing to detect a gene that was detected in samples enriched using only the poly(A)-enrichment or rRNA-, poly(A)-depletion was due to a lack of designed probes for the gene. Because the *B. malayi* annotation is currently being updated, the baits were designed with an older annotation than the one used for feature counting. Similarly, the bait design for wBm was based on the annotation published by Foster et al [18] while the annotation used for feature counting originated from GenBank (NC_006833.1). In both cases, there were genes able to be detected with the poly(A)-enrichment or depletion, but not the Agilent SureSelect. Additionally, several genes were observed to have significantly higher expression values in the poly(A)-enrichment or rRNA, poly(A)-depletion samples compared to their Agilent SureSelect counterpart.

These results illustrate the power and advantages of using probe-based enrichment to facilitate the analysis of samples that are the most biologically relevant, like those in animal tissue at a biologically relevant multiplicity of infection. The Agilent SureSelect platform serves as a method to extract reads from a secondary or tertiary organism in a sample as well as to detect low abundance transcripts. Using this platform, we successfully extracted a sufficient number of reads to represent the *B. malayi* and wBm transcriptomes in the vector portion of the *B. malayi* life cycle, the *A. fumigatus* transcriptome in a mouse infection, and the wBm transcriptome in the mammalian portion of the *B. malayi* life cycle. Provided a transcriptome experiment has an adequate bait design, we believe the Agilent SureSelect platform to be ideal and necessary in enriching samples for multi-species transcriptomic experiments, especially those in which secondary organisms are of low abundance.

## Acknowledgements

This project was funded by federal funds from the National Institute of Allergy and Infectious Diseases, National Institutes of Health, Department of Health and Human Services under grant number U19 AI110820.

## Author Contributions

MC and JCDH wrote the manuscript. LT, HL, SL, NK, XZ, and REB performed sample preparation, library construction, capture experiments, and/or sequencing. MC and AS conducted bioinformatic data analysis. LJT, LS, CMF, DAR, JMF, VMB, and JCDH were involved in study design and data interpretation. SGF and MLM provided samples. All authors read, edited, and approved the manuscript.

## Data availability

The data set(s) supporting the results of this article are available in the Sequence Read Archive (SRA) repository. The *B. malayi* datasets are available in SRP068692, the wBm datasets are available in SRP068711, and the *A. fumigatus* datasets are available in PRJNA421149.

## Supplementary Files

**Supplementary Table 1: RNA-Seq statistics for poly(A)-enriched Agilent SureSelect and poly(A)-enrichment libraries from *B. malayi* samples collected following infection of the mosquito vector**

**Supplementary Table 2: RNA-Seq statistics for Agilent SureSelect and poly(A)-enrichment libraries from *A. fumigatus***

**Supplementary Table 3: RNA-Seq statistics for Agilent SureSelect and rRNA-, poly(A)-depletion libraries from *w*Bm**

**Supplementary Table 4: RNA-Seq statistics for **B. malayi** transcriptome data using Agilent SureSelect with total RNA or poly(A)-enriched RNA**

